# *Ndufs4* knockout induces transcriptomic signatures of Alzheimer’s Diseases that are partially reversed by mitochondrial complex I inhibitor

**DOI:** 10.1101/2024.02.20.581247

**Authors:** Huanyao Gao, Kate Jensen, Jarred Nesbit, Mark Ostroot, Priyanka Baloni, Cory Funk, Eugenia Trushina

**Affiliations:** Department of Molecular Pharmacology and Experimental Therapeutics, Mayo Clinic, 200 First St. SW, Rochester, MN 55905, USA; Department of Neurology, Mayo Clinic, 200 First St. SW, Rochester, MN 55905, USA; School of Health Sciences and Purdue Institute for Integrative Neuroscience, Purdue University, West Lafayette, IN 47907, USA; Institute for Systems Biology, Seattle, WA, 98109-5263, USA

**Keywords:** mitochondrial complex I, *Ndufs4* knockout mice, transcriptomics analysis, Alzheimer’s Disease, mitochondria-targeted therapeutics, mild complex I inhibitors

## Abstract

Mitochondrial dysfunction is well documented in Alzheimer’s Disease (AD). However, whether it instigates the onset of AD remains unclear. We demonstrate that a reduction of complex I activity in wild type (WT) mice caused by a global knockout of *Ndufs4*, an accessory mitochondrial complex I subunit, was sufficient to induce transcriptomic changes in the brain reminiscent of those observed in AD patients and familial mouse models of AD. Reduced complex I activity affected expression of genes in the networks related to mitochondrial homeostasis, neuronal and synaptic function. Transcriptomic signatures in male and female *Ndufs4^-/-^*mice reflected a different severity of AD phenotype. Unexpectedly, these changes were partially rescued by a neuroprotective small molecule mild complex I inhibitor CP2. Consistent with studies in AD mice, CP2 treatment in *Ndufs4^-/-^* mice augmented the expression of genes associated with mitochondrial biogenesis and turnover, synaptic activity, autophagy, redox balance, and reduced expression of genes related to inflammation. Female *Ndufs4^-/-^* mice demonstrated a greater reversal of gene expression toward WT mice. These studies provide further support for mitochondria as a causative factor of AD pathophysiology and complex I as a putative therapeutic target.

## Introduction

Alzheimer’s Disease (AD) is the most common form of dementia with multiple processes affected years before the onset of clinical symptoms^1,2^. Traditionally, AD has been viewed as a central nervous system (CNS) disorder where the amyloid cascade hypothesis connected the accumulation of amyloid beta (Aβ) with neurodegeneration^3^. However, clearance of Aβ by anti-amyloid antibodies achieves only modest reduction in cognitive decline, with cortical atrophy persisting^4^. It is now recognized that early mechanisms of neuronal damage in AD include abnormal energy metabolism and mitochondria dysfunction^5–8^. Together with ATP production, calcium buffering and the initiation of apoptosis, mitochondria regulate cellular homeostasis, epigenetic changes, inflammation, antioxidant, and adaptive stress responses^9,10^. Most relevant to AD, mitochondrial signaling was found to be upstream of Amyloid Precursor Protein (APP) processing, Aβ production^11^, apolipoprotein E (APOE) expression^12^, and Tau pathology^13,14^. The essential role in neuronal health is further recognized by how mitochondria produce energy under sustained stress to ensure the survival of brain cells^15^ and independently affect disease progression and etiology^16–19^. The mitochondrial cascade hypothesis proposes that an individual’s genetic predisposition, environmental exposure, and lifestyle affect mitochondrial fitness and mediate and/or contribute to a variety of AD pathologies^6,18^. However, it remains unknown how mitochondria contribute to AD onset.

A prominent feature detected in AD patient brains even after accounting for neuronal loss is the reduced expression of mitochondrial proteins and RNA, particularly the enzymes and proteins involved in oxidative phosphorylation (OXPHOS)^20–24^. One of the five OXPHOS complexes, mitochondrial complex I (NADH:ubiquinone oxidoreductase), a rate-limiting enzyme in the electron transport chain, plays an essential role in cellular metabolism^25^. Mammalian complex I comprises 45 subunits^26^. Fourteen core subunits are directly involved in the catalytic reactions and are evolutionarily conserved. Seven of these core subunits are encoded in the mitochondrial DNA (mtDNA). The remaining accessory subunits are encoded in the autosomal and sex chromosomes and are required for assembly and stability of the complex^26^. One such accessory subunit is the 18 kDa NADH Dehydrogenase (Ubiquinone) Fe-S protein 4 (Ndufs4). Mutations in the *NDUFS4* gene in humans cause mitochondrial disease known as Leigh syndrome, an early-onset neurological disorder^27^. Recent development of a mouse with global *Ndufs4* knockout (*Ndufs4^-/-^*) provided a mechanistically relevant model to test strategies for this incurable disease^27–29^. Homozygous *Ndufs4*^-/-^ mice displayed increased inflammation in brain and periphery, neurodegeneration, microgliosis, abnormal lipid homeostasis, increased production of reactive oxygen species (ROS), mitochondrial dysfunction, progressive encephalopathy, and death around two months of age^27,28^. Global homozygous deletion of *Ndufs4* in mice led to tissue-specific partial loss of complex I activity with remaining ∼26% of activity in the brain tissue^30^. Metabolomics analysis in the brain tissue of *Ndufs4^-/-^* mice identified disrupted redox balance, increased glycolysis, altered fatty acid β-oxidation and amino acid homeostasis, and significant reduction in components of the tricarboxylic acid (TCA) cycle. Remarkably, similar abnormalities were reported in both AD patients and multiple mouse models of AD^31–33^. Together with abnormal OXPHOS and mitochondrial dysfunction, *Ndufs4^-/-^* mice have other AD deficiencies, including increased levels of Aβ_1-40_ in brain extracts^34^, disrupted neuronal transport, and decreased hippocampal neurotransmitters^35,36^. Strategies that reduced oxidative stress, increased mitochondrial biogenesis, levels of NAD^+^ and α-ketoglutarate, or inhibited the mammalian target of rapamycin (mTOR) and toll-like receptor 4 signaling increased *Ndufs4^-/-^* life span^27^.

Previously, we identified the small molecule tricyclic pyrone compound CP2 as a mild mitochondrial complex I inhibitor^37–43^. Continuous CP2 treatment in multiple mouse models of AD, starting at *in utero*, pre- or symptomatic stages, reduced Aβ and pTau accumulation, inflammation, oxidative damage, and improved brain and peripheral energy homeostasis. Enhanced mitochondrial bioenergetics and restored axonal trafficking helped prevent neurodegeneration and cognitive dysfunction^40,43^. In aged mice, CP2 treatment reduced levels of senescent cells and improved healthspan^40^. RNA-seq data generated in brains of CP2-treated APP/PS1 mice and cross-validated with a human postmortem dataset from the Accelerating Medicines Partnership in AD (AMP-AD) demonstrated restored gene expression in pathways most important to human disease, including redox, inflammation, and synaptic function^40^. We linked the mechanism of action to the induction of a mild energetic stress and activation of multifaceted adaptive stress responses similar to those observed in caloric restriction and exercise, likely through the AMPK pathway^39^. However, it remains unclear if CP2 actions are exclusively mediated through its binding to complex I.

The objective of this work was to determine whether loss of complex I activity in *Ndufs4^-/-^* mice abrogates the ability of CP2 to activate neuroprotective mechanisms. We conducted RNA-seq expression analysis in brain tissue from male and female *Ndufs4^-/-^* mice treated with vehicle or CP2. We found that the reduction of complex I activity in *Ndufs4^-/-^* mice was sufficient to induce transcriptomic signatures observed in AD patients and AD mice^44^. These signatures were partially rescued following CP2 treatment.

## Results

### *Ndufs4* knockout affects expression of genes involved in mitochondrial homeostasis

The *Ndufs4* whole-body knockout mice contain a deletion of exon 2, resulting in a frameshift that prevents Ndufs4 protein formation^28^. While previous metabolic and proteomic studies identified pathways affected in various tissues of *Ndufs4*^-/-^ mice^35,45^, transcriptomic data in the brain were generated only in the cerebellum without sex stratification^46^. We performed RNA-seq analysis in cortico-hippocampal tissue of 40 – 48-days-old male and female *Ndufs4^-/-^* and WT mice. All *Ndufs4^-/-^*mice were authenticated via PCR. Complete loss of exon 2 of *Ndufs4* gene was confirmed using RNA-seq data (Supplementary Fig. 1). The differential gene expression analysis between *Ndufs4^-/-^* and WT mice was done using a cutoff *p*-value < 0.05 and a log2 fold change > 0.25 (Fig. 1, Supplementary Table 1). The number of differentially expressed genes (DEGs) was relatively similar in *Ndufs4^-/-^*males and females (Fig. 1a,b). The DEGs in male and female mice had limited overlap of 190 (∼22%) upregulated and 224 (∼26%) downregulated genes, respectively (Fig. 1c). Pathway analysis revealed that mitochondrial homeostasis and fatty acid metabolism were among the top downregulated pathways shared by male and female *Ndufs4^-/-^* mice. Although not identical, pathways related to neuronal system/synapses were also downregulated in *Ndufs4^-/-^* males and females (Supplementary Fig. 2a,c). Sex-specific differences were observed in pathways enriched for upregulated genes (Supplementary Fig. 2b,d). In females, upregulated genes were enriched in TNF-α/NF-κB and BDNF signaling, DNA repair, and hypoxia-induced signaling pathways. In males, upregulated genes were primarily enriched in neuronal/synapse pathways such as neuron projection, neurotransmitter receptor activity, postsynaptic density, and modulation of synapse pathways. While pathways involved in neuronal system/synapses were downregulated in male and female *Ndufs4^-/-^* mice (Supplementary Fig. 2a,c), significant number of genes involved in synapse-related pathways were upregulated only in male mice (Supplementary Fig. 2b,d). These findings imply activation of sex-specific mechanisms in response to reduced complex I activity in *Ndufs4^-/-^*mice.

**Fig. 1:**
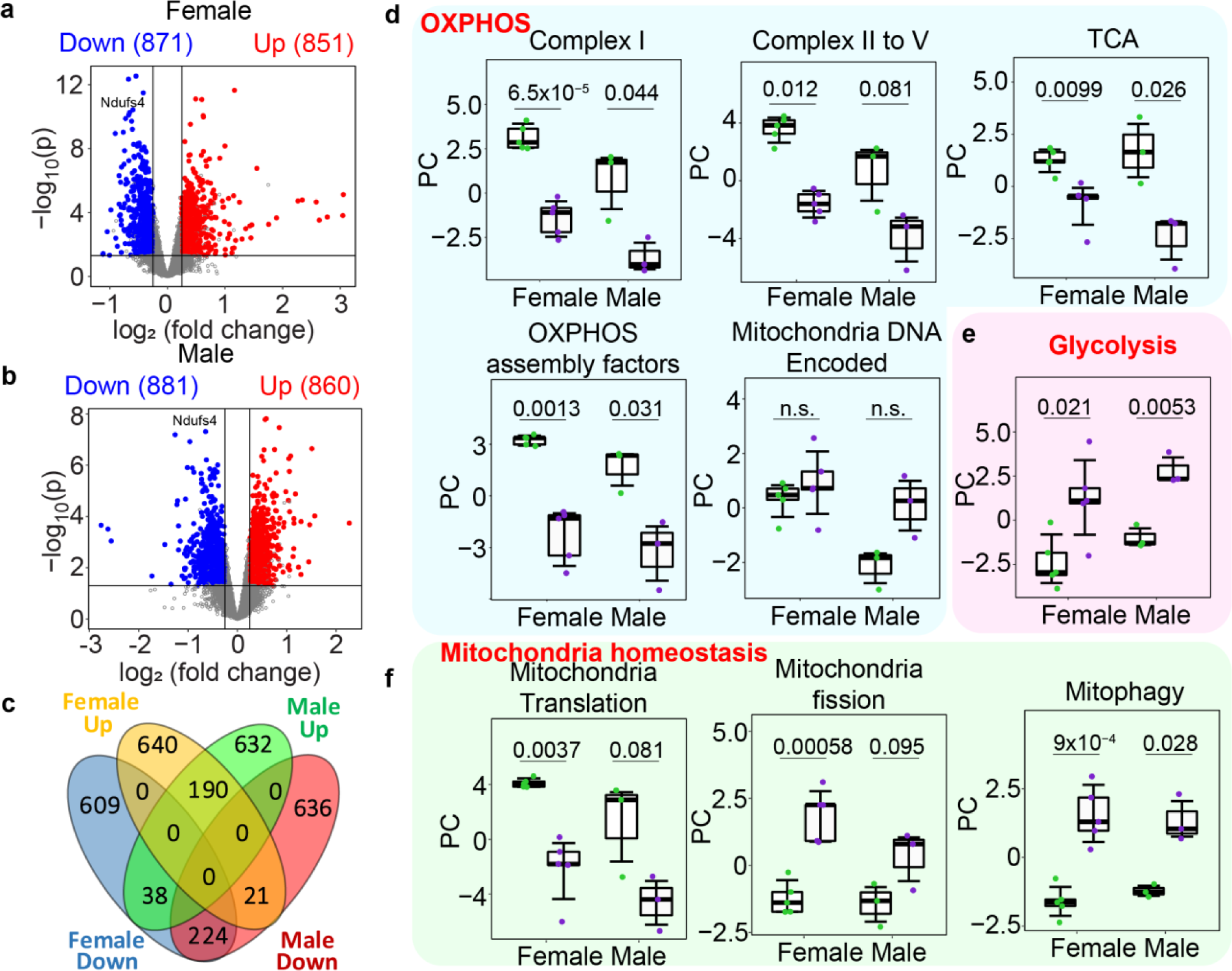
Differentially expressed genes (DEGs) in cortico-hippocampal tissue of male and female *Ndufs4^-/-^*vs WT mice. **a,b** Volcano plots showing DEGs for (**a**) female and (**b**) male mice. The x-axis represents log2 fold-change between *Ndufs4^-/-^* and WT mice, the y-axis represents negative log10 *p*-values. The number of significantly changed genes is shown on top. Red, upregulated genes; blue, downregulated genes. **c** A Venn diagram showing the overlap between up- and downregulated DEGs for female and male mice. **d–f** The first principal component (PC) plot of sample-wise Z-score transformed expressions of all genes (mtDNA encoded) or DEGs for all other pathways in OXPHOS and mitochondrial homeostasis. WT, green; *Ndufs4^-/-^*, purple. The directionality of PCs was adjusted to reflect the expression level. Each dot represents one sample. Statistical difference between groups was tested by two-tailed Student’s *t*-test with *p*-values shown in the plots. Boxes represent the interquartile range (IQR) and whiskers represent 1.5 × IQR. *N* = 3 for *Ndufs4^-/-^* and WT males, *n* = 5 for *Ndufs4^-/-^* and WT females. OXPHOS: oxidative phosphorylation; TCA: tricarboxylic acid cycle.

We next examined the expression of genes involved in the OXPHOS and mitochondrial homeostasis. To quantitatively compare the integrated expression of all DEGs of a specific pathway, we calculated the first principal component (PC) of DEGs (Fig. 1d-f). The expression of all OXPHOS complexes was reduced in *Ndufs4^-/-^*males and females (Fig. 1d) despite not reaching significance in the expression of genes of complexes II - V in males, likely due to a variance among samples (Fig. 1d). The expression of genes involved in the TCA cycle (e.g., *Bckdhb* and *Mrps36* related to α-ketoglutarate, *Sfxn5* related to citrate transporter, and *Pcx* and *Pck2* involved in the conversion of pyruvate to oxaloacetate) as well as OXPHOS complex assembly factors was significantly reduced in male and female *Ndufs4^-/-^* mice (Fig. 1d, Supplementary Table 1). We calculated PCs for all mtDNA encoded genes and found no significant changes, which was not surprising, given that *mt-nd2* was the only significantly differentially expressed mtDNA gene (Fig. 1d, Supplementary Fig. 3). In *Ndufs4^-/-^* males and females, genes involved in glycolysis were significantly upregulated, consistent with the activation of an alternative ATP-producing pathway (Fig. 1e). Mitochondrial translation was significantly downregulated while genes involved in mitochondrial fission and mitophagy were significantly upregulated, indicating a disturbance of mitochondrial homeostasis (Fig. 1f). Therefore, loss of complex I activity causes comprehensive alterations in genes involved in mitochondrial dynamics and function in male and female *Ndufs4^-/-^* mice.

### CP2 treatment in *Ndufs4^-/-^* mice modulates sex-specific expression of genes involved in OXPHOS and mitochondrial homeostasis

We previously demonstrated that CP2, a mild complex I inhibitor, enhances mitochondrial biogenesis, dynamics, and function, increasing levels of deacetylases Sirtuins 1 and 3, and reduces oxidative damage in AD mice^40,41,43^. Since strategies that reduce ROS and improve mitochondrial function have been shown to enhance life span in *Ndufs4^-/-^* mice^27^, we administered CP2 to *Ndufs4^-/-^* males and females via drinking water *ad lib* (25 mg/kg/day) starting at 21 days through life (Fig. 2a). While CP2 treatment did not induce side effects and did not impact weight gain demonstrating the lack of toxicity (Supplementary Fig. 4), it also did not affect lifespan (Fig. 2a).

**Fig. 2:**
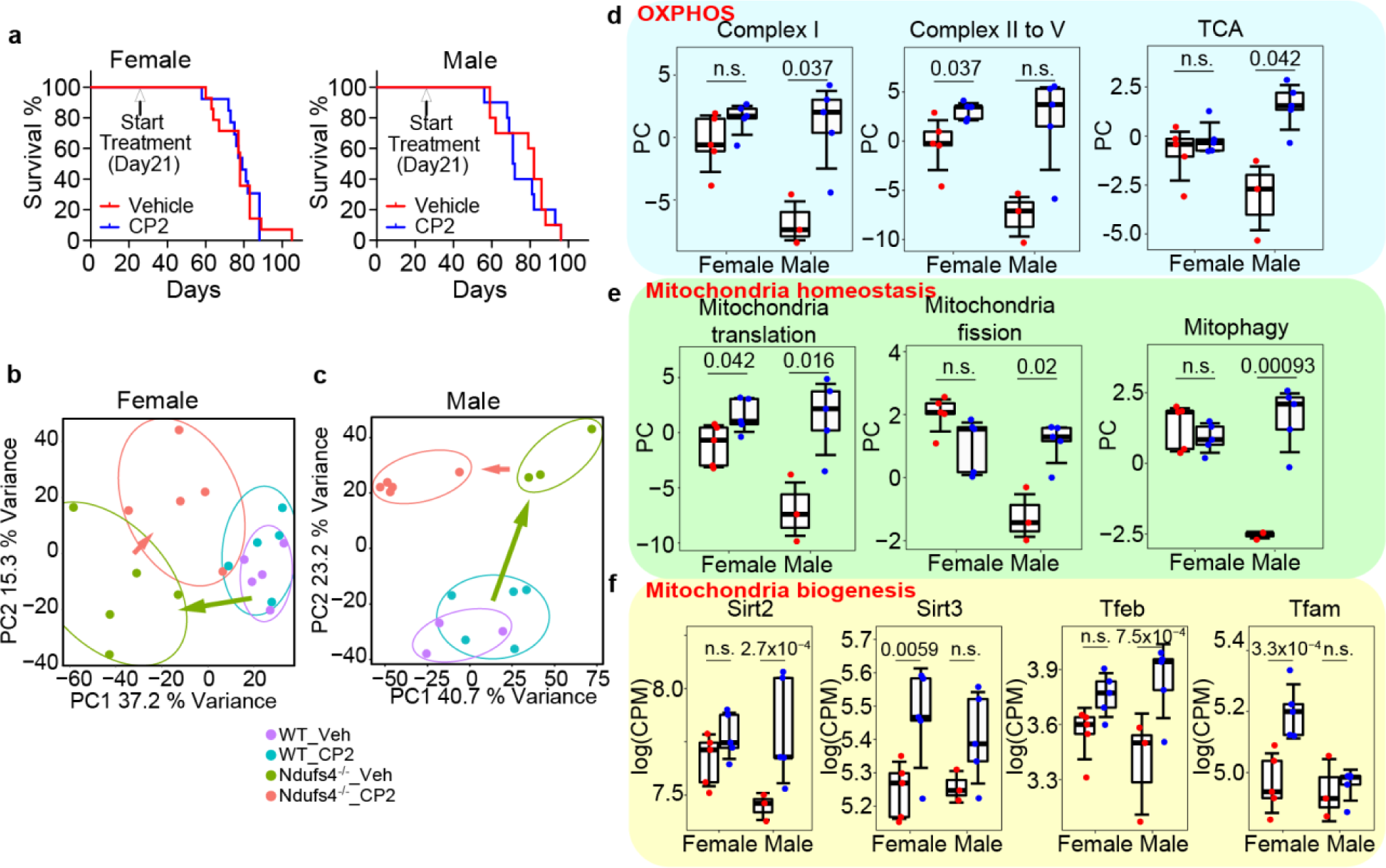
CP2 treatment reverses gene expression profile in brain tissue of *Ndufs4^-/-^* mice. **a** Survival curves of male and female *Ndufs4^-/-^* mice treated with vehicle (red) or CP2 (blue). *N* = 14 for females and males treated with vehicle; *n* = 13 for females and males treated with CP2. **b,c** The PCA plots of DEGs for (**b**) female and (**c**) male *Ndufs4^-/-^* and WT mice treated with vehicle or CP2. **d-e** The first PC of sample-wise Z-score transformed expression of DEGs between CP2 and vehicle treated *Ndufs4^-/-^* mice for specific pathways. Red, vehicle; blue, CP2. The directionality of PCs was adjusted to reflect the expression level. Each dot represents one sample. Statistical difference between groups was tested using a two-tailed Student’s *t*-test with *p*-values marked in the plots. **f** The expression of mitochondrial biogenesis genes in CP2 and vehicle treated *Ndufs4^-/-^* mice. Each dot represents one sample. Red, vehicle; blue, CP2. Statistical difference between groups was tested by GLM model using EdgeR; the *p*-values are marked in the plots. Boxes in boxplots represent IQR and whiskers represent 1.5 × IQR. *N* = 3 for vehicle treated *Ndufs4^-/-^*and WT male groups, *n* = 5 for CP2 treated *Ndufs4^-/-^* and WT male groups; *n* = 5 for all female groups. CPM: counts per million; OXPHOS: oxidative phosphorylation; PC: principal component; PCA: principal component analysis; TCA: tricarboxylic acid cycle.

We therefore tested whether reduced complex I assembly in *Ndufs4^-/-^*mice abrogated CP2’s neuroprotective mechanisms, which could explain the lack of its effect on life extension. Male and female WT and *Ndufs4^-/-^*mice 37 - 45-days-old were gavaged with vehicle or CP2 (25 mg/kg/day) once a day for 3 consecutive days. At the end of treatment, cortico-hippocampal tissues were subjected to the RNA-seq analysis (Fig. 2b-f), and differential expression analysis was performed using the same cutoff as for WT and *Ndufs4^-/-^* mice. The analysis revealed significantly greater number of altered genes in CP2 treated males (1312 upregulated/1049 downregulated) compared to females (457 upregulated/ 431 downregulated) with only 92 upregulated/33 downregulated overlapping genes between sexes (Supplementary Fig. 5a-e, Supplementary Table 2). In male and female *Ndufs4^-/-^* mice, CP2 treatment upregulated mitochondrial translation and biogenesis, evident by increased expression of critical transcription factors (Fig. 2e,f). The expression of genes involved in OXPHOS subunits, assembly factors, and mitochondrial homeostasis was better restored in *Ndufs4^-/-^* males. Specifically, significant increase was observed in the expression of genes related to integrated complex I, the TCA cycle, OXPHOS assembly factors, mitochondrial fission and mitophagy (Fig. 2d,e, Supplementary Fig. 5f-h). Among mtDNA-encoded genes, the expression of three out of five OXPHOS complexes was increased in CP2 treated males (Supplementary Fig. 5h). In contrast, significant increase in integrated OXPHOS complexes II – V but not complex I (Fig. 2d), as well as mtDNA-encoded complex V subunit *mt-Atp6,* was observed in CP2 treated *Ndufs4^-/-^* females. The enhanced expression of genes involved in OXPHOS and mitochondrial homeostasis in male *Ndufs4^-/-^* mice did not lead to a better transcriptome-wide reversal compared to females (Fig. 2b,c). After CP2 treatment, *Ndufs4^-/-^* females resembled the expression profile of WT females as shown in the principal component analysis (PCA) performed using all DEGs (Fig. 2b, pink arrow). CP2 treated *Ndufs4^-/-^* males had unique signatures that did not revert to the signatures of WT males (Fig. 2c, pink arrow). To quantify the differences between genotype/treatment groups, we calculated the Mahalanobis distance (MD) between PCA centroids. CP2 treatment resulted in MD change of 1.53 for *Ndufs4^-/-^* but only 0.64 for WT females. Moreover, CP2 reversed the MD from 4.75 to 3.40 between *Ndufs4^-/-^* and WT female mice (Fig. 2b). In males, although CP2 resulted in MD change of 29.6 in *Ndufs4^-/-^* mice, MD between *Ndufs4^-/-^*and WT remained almost unchanged (11.4 vs 11.2, Fig. 2c). Sex-specific differences in response to treatment were particularly evident in the pathway analysis of DEGs between CP2 and vehicle treated *Ndufs4^-/-^* mice (Supplementary Fig. 6). CP2 treatment suppressed TNFα signaling and hypoxia that were upregulated in *Ndufs4^-/-^*females, and upregulated neuronal and synaptic functions, such as glutamatergic synapse and calcium signaling (Supplementary Fig. 2a,b, 6a,b). In *Ndufs4^-/-^* males, the major pathways affected by CP2 almost exclusively related to the mRNA processing and protein translation (Supplementary Fig. 2c,d, 6c,d).

### WGCNA coexpression analysis identified specific modules rescued by CP2 treatment

The direct pathway analysis, while providing valuable information on top pathways enriched among all DEGs, is limited to a small subset of genes repetitively enriched in a few similar or related pathways, as shown in Supplementary Fig. 2 and 6. To systematically characterize the impact of *Ndufs4* knockout and CP2 treatment based on all DEGs, as well as to quantify changes in specific pathways and compare across groups, we conducted the Weighted Gene Co-Expression Network Analysis (WGCNA) using DEGs from the RNA-seq data of CP2 or vehicle treated male and female *Ndufs4^-/-^* and WT mice. Using a total of 4740 DEGs, we built 17 coexpression modules (Fig. 3a, Supplementary Fig. 7) with the most enriched pathways for each module shown in Fig. 3a (Supplementary Table 3). We tested whether the module expression was significantly different as a result of *Ndufs4* knockout or CP2 treatment by calculating the eigengene values for each module and comparing across groups (Fig. 3, Supplementary Fig. 8). The heatmaps (Fig. 3b,c) present the normalized expression for all significantly changed modules in male and female WT and *Ndufs4^-/-^*mice treated with vehicle or CP2. Modules consistently changed in male and female *Ndufs4^-/-^*mice compared to WT included the downregulated module brown (mTOR-signaling/lipid metabolism) and the upregulated module darkturquoise (mRNA processing) (Fig. 3a). Changes in module darkturquoise were reversed by CP2 treatment only in males. Changes in module brown were not affected by CP2 treatment neither in *Ndufs4^-/-^*males nor females. Distinct downregulated modules associated with neuronal and synaptic dysfunction in female *Ndufs4^-/-^* mice included modules orange (post-synapsis) and grey60 (glutamatergic synapse/Ephrin Receptor (EPHR) signaling) were CP2 treatment reversed the expression of genes in module grey60 (Fig. 3a,b). In *Ndufs4^-/-^* males, the downregulated modules included greenyellow (dopaminergic neurogenesis) and lightgreen (axon guidance), and both were rescued by CP2 treatment (Fig. 3a,c, Supplementary Fig. 8). The number of modules significantly affected by CP2 was greater in males compared to females (Fig. 3a-c), with only few modules where CP2 reverted changes induced by *Ndufs4* knockout (Fig. 3a-c). Among these were modules greenyellow and grey60 related to the improvement of synaptic function, suggesting that the reduced complex I activity in *Ndufs4^-/-^* mice did not block the ability of CP2 to activate neuroprotective mechanisms.

**Fig. 3:**
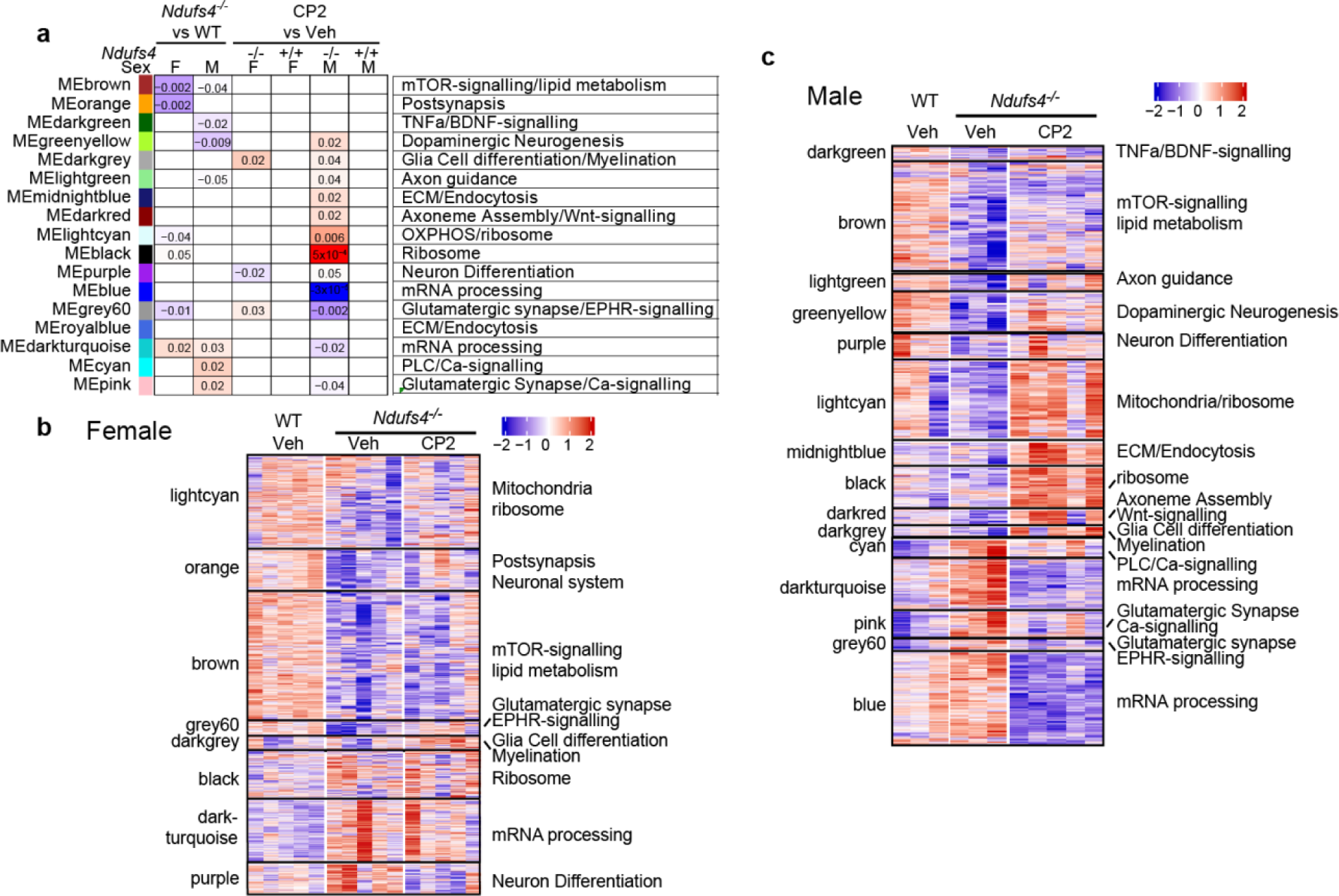
WGCNA coexpression analysis using DEGs of *Ndufs4^-/-^*mice. **A** 17 coexpression modules built using WGCNA program. Module names and colors were labeled on the left, primary pathways of genes in each module are labeled on the right. Association between different groups was compared by calculating eigengenes of each sample. Statistical difference between groups was tested using two-tailed Student’s *t*-test with *p*-values labeled and colored as heatmaps (red, upregulation; blue, downregulation). Comparison pairs are labeled on the top. **b-c** Heatmaps for modules significantly different between at least one group of comparison as shown in (**a)** for (**b**) female and (**c**) male mice, respectively. Each row represents a gene, and each column represents a sample. Module labels (left) and primary pathways (right) are as shown for each module. Expression of each gene is Z-score normalized across all samples.

### CP2 treatment reversed AD-like transcriptomic signatures in the brain of *Ndufs4^-/-^* mice

A recent meta-analysis using RNA-seq data from 2,114 post-mortem brain samples of AD and control subjects from AMP-AD cohort identified five highly conserved Clusters A-E, comprised of 30 coexpression modules across different brain regions^44^. These human coexpression modules were compared to 251 DEG sets from RNA-seq data corresponding to 96 distinct mouse studies selected for their relevance to known AD pathology, including several Aβ (APP) and pTau (the microtubule-associated protein tau (MAPT)) models. The authors found directionality concordant overlap between several mouse expression signatures and the 30 human AD-associated brain consensus coexpression modules within consensus Clusters, particularly Clusters B and C (Fig. 4a). Downregulation of Cluster C corresponded to early-stage mild symptoms, including synaptic and neuronal dysfunction, while upregulation of Cluster B corresponded to more advanced pathologies related to tau hyperphosphorylation, brain atrophy, and microgliosis. Concurrent changes in Clusters B and C were relatively rare among all models but were observed in select APP and MAPT models in a time and sex-dependent manner. The overlap with Clusters A, D and E was relatively sparse, containing more divergent pathway enrichment (Fig. 4a)^44^. We used this cross-species resource to determine to what extent transcriptomic signatures in the brain of *Ndufs4^-/-^* mice overlap with the specific human-mouse AD modules and whether CP2 treatment reversed gene expression in these Clusters (Fig. 4).

**Fig. 4:**
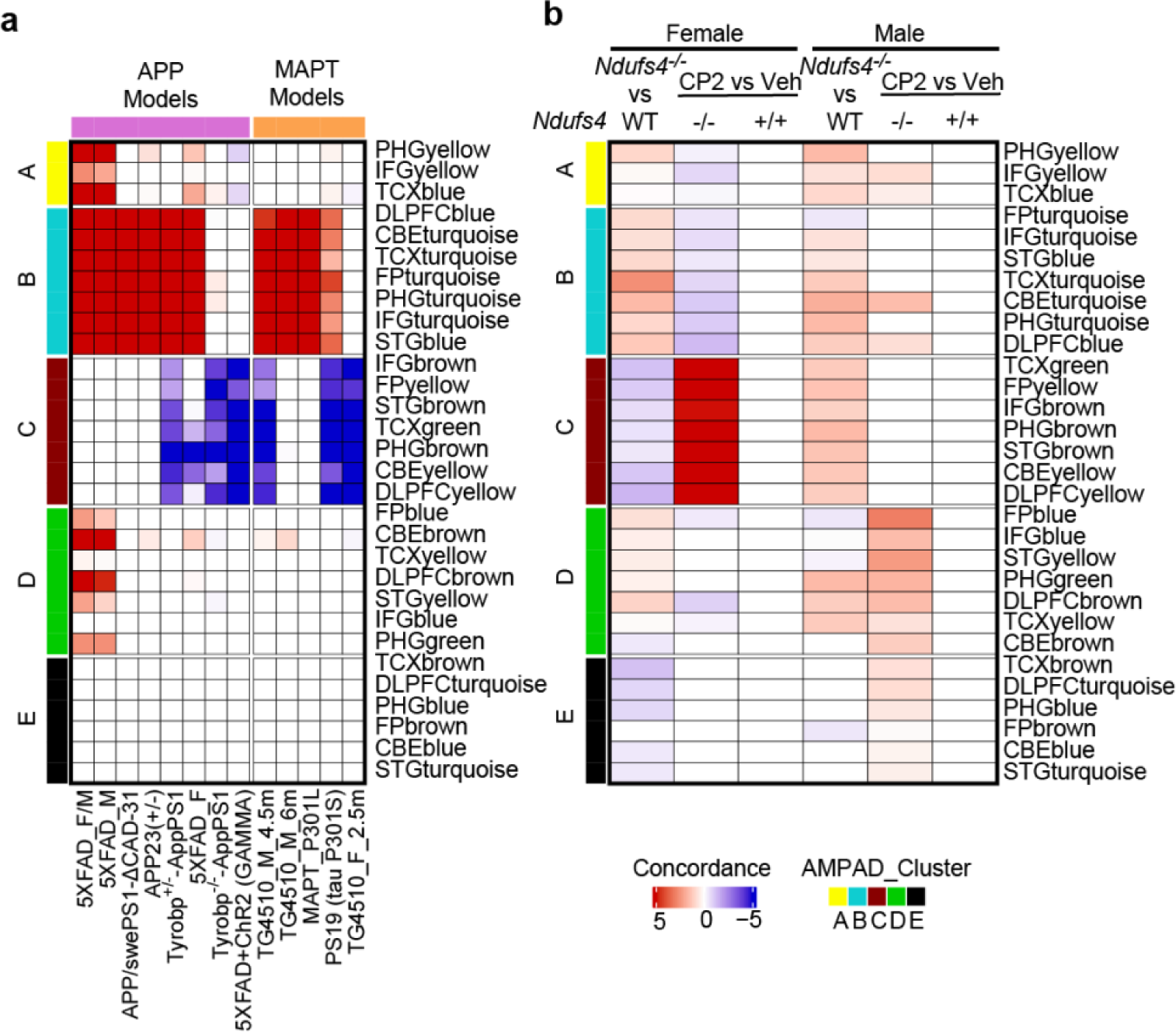
The overlap between AMP-AD aggregated coexpression modules and DEGs in CP2 or vehicle treated *Ndufs4^-/-^* and WT mice. **a** Concordance overlap between AMP-AD aggregated coexpression modules and DEGs from representative AD mouse models, as labeled on the bottom. Color bar on the left: AMP-AD aggregated coexpression clusters. Labels on the right: AMP-AD aggregated coexpression modules. Color bar on top: Pink (humanized APP/PS1 transgenic mice); orange (MAPT knock in mouse models). A heatmap color darkness represents log10 enrichment false-discovery rate adjusted *p*-values: red, concordance is more significant in positive direction; blue, concordance is more significant in negative direction. **b** An overlap concordance between AMP-AD aggregated coexpression modules and DEGs from *Ndufs4^-/-^* mice. AMP-AD Clusters (color bar on the left) and modules (labels on the right), and concordance color scale are the same as in (**a**). DEG analysis pairs for each column are labeled on top. The concordance was calculated for *Ndufs4^-/-^* vs WT columns and discordance was calculated for CP2 vs vehicle treatment (see Methods).

We found that *Nudfs4^-/-^* female mice recapitulated concordant changes throughout consensus Clusters B and C, associated with moderate to severe AD-like phenotype present in APP/PS1 model Tyrobp-KO/AppPS1, 5XFAD female mice (5XFAD_F), tau knock-in MAPT models TG4510 4.5-month-old males (TG4510_M_4.5m) and PS19 (Fig. 4a,b). The *Ndufs4^-/-^* male mice recapitulated signatures in Cluster B, and lesser in Custers A and D, but not in Cluster C, resembling more severe phenotype of AD mouse models 5XFAD mixed sex (5XFAD_F/M) or male mice only (5XFAD_M), and to a lesser extent APP/PS1, APP23(+/-),TG4510 male mice (TG4510_M_6m), and MAPT P301L. Although DEGs in *Ndufs4^-/-^* males matched well with Cluster C genes, the majority of matching appears to be upregulated instead of being downregulated as seen in the AD mouse models. CP2 treatment reversed pathological expression profile in Clusters B and especially in Cluster C in *Ndufs4^-/-^* female mice while treatment had no reversal effect on gene expression in either Clusters B or C in males (Fig. 4b). Interestingly, male *Ndufs4^-/-^* mice recapitulated similar increase in gene expression in Cluster B as female mice. However, CP2 treatment did not revert these signatures in males.

We next took a closer look at DEGs matched between *Ndufs4^-/-^*mice and AMP-AD patient Clusters. The genes concordantly upregulated in Cluster B for AD patients and female *Ndufs4^-/-^*mice were enriched in pathways related to phagocytosis, adipogenesis, vascular system, including transport across the blood-brain barrier (Fig. 5a,b,d). The upregulation of complement activation pathways was shared by male and female AD patients and *Ndufs4^-/-^* mice, consistent with the overall cell type enrichment of microglia, pericyte, and endothelial cells indicative of increased inflammation in AD^44^ (Fig. 5d,e). Genes concordantly downregulated in Cluster C for female AD patients and *Ndufs4^-/-^* mice were enriched in neuronal system and synapses pathways (Fig. 5g,i), also consistent with the overall cell type enrichment of neuron and early-stage synaptic and neuronal dysfunction as described in the original publication^44^. CP2 treatment reversed the concordant genes in Clusters B and C in females but not in male mice (Fig. 5c,h), despite that the enriched pathways were comparable to female mice. Particularly, DEGs in male mice had elevated expression in Clusters B and C concordantly with male patients, but CP2 further elevated these genes in both Clusters (Fig. 5c,h).

**Fig. 5:**
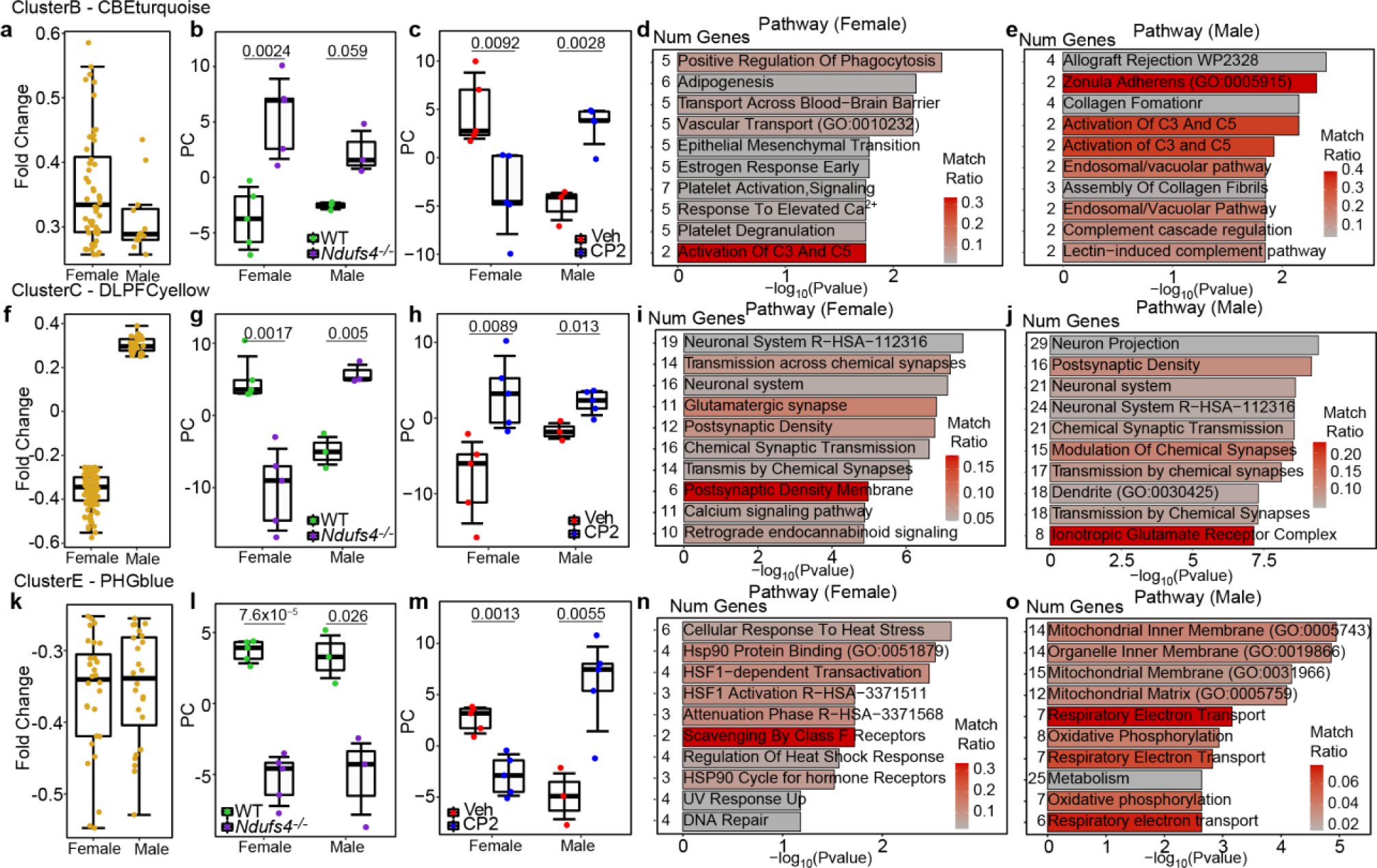
Analysis of the overlap between AMP-AD modules and DEGs of *Ndufs4^-/-^* mice. **a, f, k** Fold change in expression of DEGs between AD and Normal samples in AMP-AD (**a**) CBEturquoise, (**f**) DLPFCyellow, and (**k**) PHGblue modules that overlap concordantly with DEGs of *Ndufs4^-/-^* vs WT male and female mice. Each dot represents a gene. **b, g, l** First PC of sample-wise Z-score transformed expression of DEGs of *Ndufs4^-/-^* vs WT mice concordantly overlaps with AMP-AD (**b**) CBEturquoise, (**g**) DLPFCyellow, and (**l**) PHGblue modules, colored by mouse genotype (WT, green; *Ndufs4^-/-^*, purple) and grouped by sex. Directionality of PCs was adjusted to reflect the expression level. **c, h, m** First PC of sample-wise Z-score transformed expression of DEGs of CP2 vs vehicle treated *Ndufs4^-/-^* mice that discordantly overlapped with AMP-AD (**c**) CBEturquoise, (**g**) DLPFCyellow, or (**k**) PHGblue modules, colored by treatment (vehicle, red; CP2, blue) and grouped by sex. The directionality of PCs was adjusted to reflect the expression level. For all boxplots, each dot represents one sample. Statistical difference between groups was tested using two tailed Student’s *t*-test with *p*-values marked in the plots. Boxes represented IQR and whiskers represented 1.5 × IQR. *N* = 3 for vehicle treated *Ndufs4^-/-^*males. *N* = 5 for all other groups. **d, e, I, j, n, o** Pathway analysis of overlapped genes between DEGs of AMP-AD (**d**) CBEturquoise module for females, (**e**) CBEturquoise module for males, (**I**) DLPFCyellow module for females, (**j**) DLPFCyellow module for males, (**n**) PHGblue module for females, (**o**) PHGblue module for males, and *Ndufs4^-/-^* vs WT mouse DEGs (concordantly) or CP2 vs vehicle mouse DEGs (discordantly) using Enrichr database. X axis represents negative log *p*-values of pathway enrichment, and color ramp represents percentage of genes matched for a specific pathway. PC: principal component.

Mitochondria-related genes were mostly enriched in Cluster E. Both *Ndufs4^-/-^* females and males had moderate concordantly downregulated genes overlapped with Cluster E, but only in males was there an enrichment in mitochondrial genes (Fig. 5k,l,n,o). CP2 treatment increased gene expression in Cluster E only in male *Ndufs4^-/-^* mice (Fig. 5m), implicating better reversal of mitochondrial homeostasis, as we have demonstrated earlier (Fig. 2d,e, 3a-module lightcyan). In summary, the concordance analysis between AD patients and *Ndufs4^-/-^* DEGs supports our findings that although gene expression related to OXPHOS and mitochondrial homeostasis was better reversed by CP2 in male mice, females had a better reversal of signatures associated with the early (Cluster C) and late (Cluster B) disease stages, particularly in terms of reduction of inflammation and recovery of pathways related to neuronal function. This may be associated with the presence of early-stage phenotype as represented by Cluster C in female mice but not male mice, indicating the potential importance of early intervention.

Finally, we examined whether the overlapped DEGs between *Ndufs4^-/-^* mice and AMP-AD Clusters A-E were specifically enriched in certain coexpression modules built from male and female *Ndufs4^-/-^*DEGs combined together. We found that Cluster B genes were most enriched in five *Ndufs4^-/-^* modules: midnightblue and royalblue (extracellular matrix/endocytosis), black (ribosome), darkgreen (TNFα/BDNF signaling), and darkred (axoneme assembly/Wnt-signaling) (Fig. 6a). While most of the modules overlapped with several upregulated human DEGs in Cluster B, only the black module showed consistent and significant upregulation in female knockout mice (Fig. 6b). This module was not rescued by CP2 treatment (Fig. 6b, 3b). Cluster C genes were enriched in six *Ndufs4^-/-^* modules: grey60 (glutamatergic synapse/EPHR signaling), orange (post-synapsis/neuronal system), greenyellow (dopaminergic neurogenesis), pink (glutamatergic synapse/Calcium signaling), purple (neuron differentiation), and cyan (phospholipase C/Calcium signaling) (Fig. 6a). Essentially, all modules were related to neuronal functions and development.

**Fig. 6:**
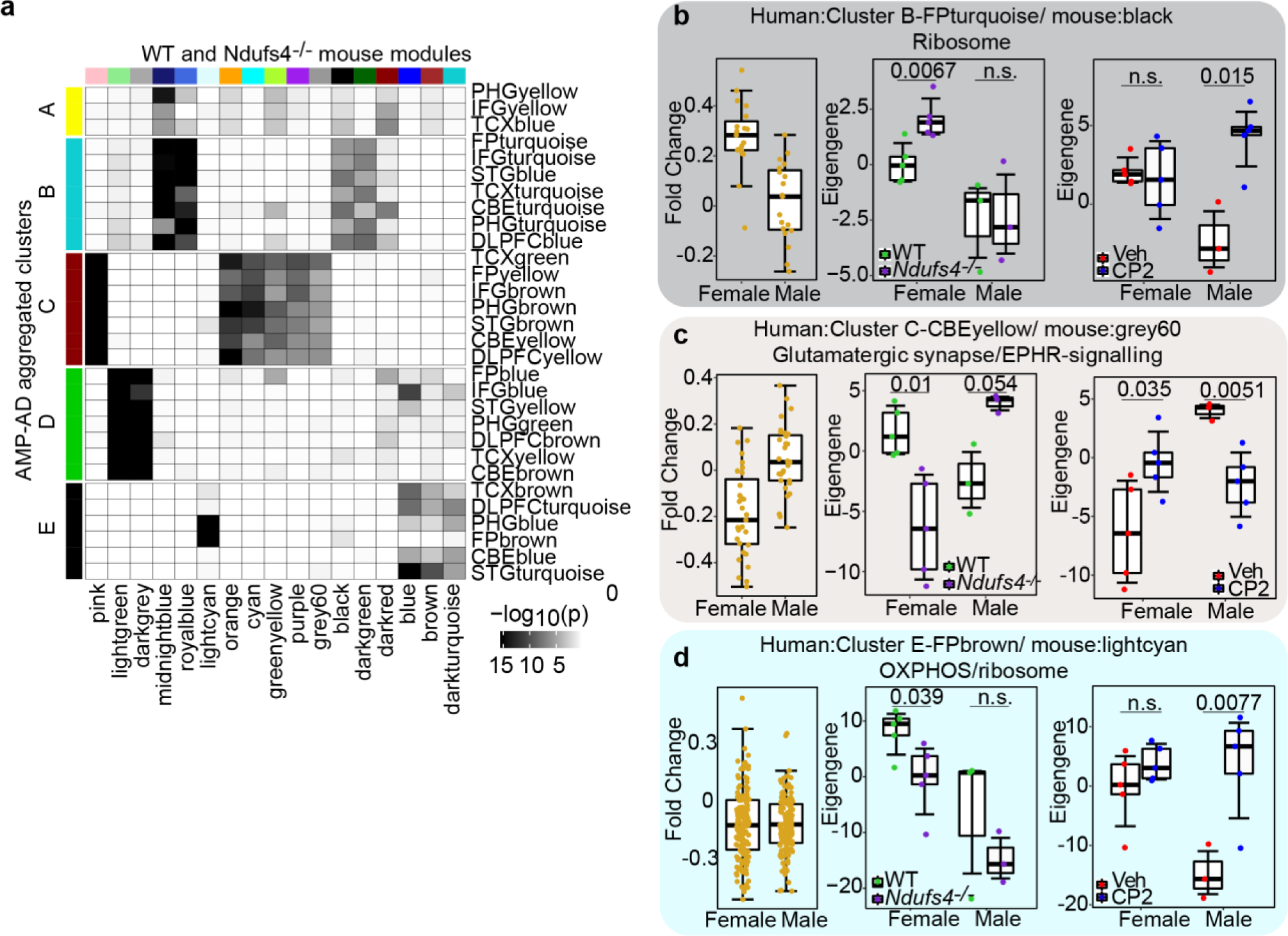
Overlap between AMP-AD aggregated coexpression modules and WGCNA coexpression modules built by DEGs of *Ndufs4^-/-^* mice. **a** The overlap enrichment (hypergeometric test) AMP-AD Clusters (color bar on the left) and modules (labels on the right), and WGCNA coexpression modules built from *Ndufs4^-/-^* mouse models as labeled at the bottom and color coded on top. Enrichment was in grey scale by log10 (*p*-values). **b-d** Examples of concordantly overlapped genes between AMP-AD and mouse modules. For each overlapped module-pairs, left: overlapped genes in corresponding AMP-AD module, as labeled on top of the sub-figure; middle: eigengene plot comparing vehicle treated *Ndufs4^-/-^*and WT groups, recalculated using AMP-AD / *Ndufs4^-/-^* module overlapped genes; right: eigengene plot comparing CP2 and vehicle treated *Ndufs4^-/-^* groups, recalculated using AMP-AD / *Ndufs4^-/-^* module overlapped genes. All eigengene plots were statistically tested by unpaired two-tailed Student’s *t*-test. Boxes represented IQR and whiskers represented 1.5 × IQR. *N* = 3 for vehicle treated *Ndufs4^-/-^* males. *N* = 5 for all other groups.

Genes in module grey60 (glutamatergic synapse/EPHR signaling) were significantly downregulated in female AD patients and *Ndufs4^-/-^* female mice. CP2 treatment rescued gene expression in this module, consistent with the whole module eigengene analysis (Fig. 6c, 3b). Although changes in multiple modules were observed in *Ndufs4^-/-^* male mice, they did not correlate with changes in AMP-AD DEGs of AD male patients. Another interesting finding was related to module lightcyan. This module was enriched in OXPHOS/ribosome genes where overlapped human-mouse DEGs indicated moderate downregulation (Fig. 6d). The downregulation was significant only for female *Ndufs4^-/-^* mice with the downtrend for males. CP2 treatment eliminated changes in gene expression observed in females and significantly increased the expression in males (Fig. 6d), consistent with our findings that CP2 rescued OXPHOS deficiency better in male mice but reversed the overall gene expression to that observed in WT only in female mice.

## Discussion

Several key findings resulted from this study. We have shown that knockout of complex I subunit with concomitant loss of complex I assembly and activity resulted in significant downregulation of genes involved not only in OXPHOS but also in mitochondrial dynamics and homeostasis and neuronal function indicating the central role of complex I in the regulation of complex networks in the brain. A reduction of complex I activity in WT mice was sufficient to induce transcriptomic signatures in specific coexpression Clusters shared between AD patients and multiple mouse models of familial AD with Aβ and pTau pathology. These AD-like transcriptomic changes could be partially reversed by a small molecule mild complex I inhibitor. Male and female mice have differential response to the loss of complex I activity, which appears to affect the severity of AD-like signatures and a response to treatment.

*Ndufs4^-/-^* mice resemble Leigh syndrome, where mutations in *NDUFS4* cause the most severe clinical phenotype compared to other complex I mutations^47^. Individuals with *NDUFS4* mutations die before 3 years of age^48^. Together with symptoms that include hypotonia, visual impairment, psychomotor arrest, and respiratory failure, magnetic resonance imaging revealed abnormalities in the brainstem, basal ganglia, and the cerebral cortex. In fibroblasts and muscle tissue, *NDUFS4* mutations reduce complex I activity up to 24% and 74%, respectively^48^. Based on a very short life span of these patients, data on the effect of *NDUFS4* mutations on the development of AD are not available.

Brain tissue of AD patients exhibit decreased mtDNA copy number, and low expression of genes and levels of proteins associated with mitochondrial respiratory chain^20–23,49^. Reduced activity of complex I in AD brain tissue was recognized as the most influential pathway involved in AD pathology^50,51^. Recently published AD GWAS studies identified *NDUFS2,* one of the catalytic subunits of mitochondrial complex I, as a putative causal gene^52,53^. Despite the fact that complex I is a rate-limiting enzyme in the OXPHOS machinery, it is unclear whether a reduction of complex I activity is sufficient to initiate the disease. The complexity is associated with a multitude of compensatory pathways available to backup, at least temporarily, failing mitochondrial ability to produce ATP, and relatively high threshold required for a significant loss of the enzyme activity. However, recent investigations provided strong evidence that the reduction of complex I activity has the most detrimental effect on the brain. Experiments conducted in mouse models with either CNS-specific or the whole-body *Ndusf4* knockout showed that the impact on complex I subunit levels was the greatest in the brain compared to other organs^30,54^, and the clinical phenotype was primarily due to brain abnormalities. The NesKO mice with a specific *Ndufs4* knockout in neurons and glia cells displayed similar phenotype as the whole-body knockout mice showing progressive neuronal deterioration, gliosis, loss of motor ability, breathing abnormalities, and death at ∼7 weeks of age^55,56^. Selective inactivation of *Ndufs4* in the dorsal brain stem vestibular nucleus (VN) induced neurodegeneration, breathing abnormalities and increased mortality, which were rescued after viral restoration of *Ndufs4* expression in VN^56^. These data suggest that alterations in complex I activity associated with age or disease could disproportionally affect brain compared to other organs.

Indeed, the examination of gene expression profiles in the cortico-hippocampal brain region of male and female *Ndufs4^-/-^* mice demonstrated that the loss of ∼70% of complex I activity due to genetic ablation of a single subunit profoundly affected the expression of genes involved in global mitochondrial homeostasis. Consistent with the previous study conducted in the similar brain region using proteomics analysis^30^, *Ndufs4* knockout affected the expression of genes involved in other complex I subunits. Furthermore, we also found significant alterations in genes related to other OXPHOS complexes, OXPHOS assembly factors, components of the TCA cycle, mitochondrial dynamics, and homeostasis. Similar to those observed in AD patients, changes related to mitochondrial function were primarily restricted to the nuclear encoded genes^24^.

Balanced mitochondrial dynamics (fission, fusion, mitophagy, biogenesis and transport) are essential for the removal of damaged organelles and production of new, healthier, organelles to ensure adequate energy support for synaptic function^57^. Significantly increased mitochondrial fission and mitophagy in *Ndufs4^-/-^* mice without increased biogenesis supports a progressive loss of mitochondria. Synaptic function critically depends on energy provided by mitochondria. Not surprisingly, our analysis identified multiple pathways related to neuronal function and synapse among the most affected in *Ndufs4^-/-^* males and females. However, the directionality of changes was not the same. In *Ndufs4^-/-^*females, these pathways were consistently downregulated while in males they were up- and downregulated. This may indicate a different trajectory of neuronal dysfunction, and the overall trajectory of the development of AD-like phenotype, in males and females, possibly due to distinct ability to activate compensatory mechanisms. Indeed, in females, among activated pathways were the ones related to hypoxia. Previous studies demonstrated that oxidative stress and increased ROS production elevated levels of O_2_ in the brain of *Ndufs4^-/-^* mice, and strategies that induced hypoxia, such as breathing carbon monoxide, rescued the disease^29,27^. Interestingly, carbon monoxide modulates cellular response to stress and adaptive signaling via inhibition of mitochondrial complex IV^58^ suggesting that additional inhibition of OXPHOS in *Ndufs4^-/-^* mice is not detrimental.

In support of the critical importance of complex I activity, we found that *Ndufs4^-/-^* mice developed AD-specific transcriptomic changes shared by AD patients and multiple mouse models of familial AD. Mechanisms recapitulated by these signatures included abnormal energy and lipid homeostasis, loss of synaptic function, inflammation, increased Aβ levels, abnormal endocytic pathway, mTOR, and loss of mitochondrial dynamics and function that are well documented in AD GWAS studies^35,36,44,46,55^. The coexpression Clusters that captured the AD-like phenotype in patients represent different stages of the disease and severity of symptoms^44^. The extent of AD-like signatures differed in *Ndufs4^-/-^* males and females. The *Ndufs4^-/-^*females recapitulated changes in clusters related to the mild AD phenotype, while *Ndufs4^-/-^* males mimicked signatures of more severe AD. These sex-specific differences modified the response to CP2 treatment. One of the counterintuitive mechanisms of improving mitochondrial function includes mild inhibition of OXPHOS, complex I in particular^39^. The mechanism of action includes the multifaceted adaptive response to mild energetic stress and activation of multiple neuroprotective mechanisms. Interestingly, as is the case of complex I inhibitor metformin or complex I, III and V inhibitor resveratrol, continuous application of these compounds is safe^39,59^. Consistent with our previous findings in AD mice^37–41,43^, 3-day CP2 treatment in *Ndufs4^-/-^* males and females enhanced mitochondrial biogenesis, evident by increased mitochondrial translation and activation of transcription factors Tfeb and Tfam, which lead to the augmented expression of genes involved in the OXPHOS, the TCA cycle, and the mitochondria specific neuroprotective deacetylase Sirtuin 3^60^. Concomitantly increased mitochondrial biogenesis, translation and mitophagy indicate augmented mitochondrial turnover and replenishment. Activation of Tfeb also suggests improved lysosomal degradation via autophagy, a process that is stalled in AD contributing to disease pathogenesis^61,62^. Despite a reversal of the expression of mitochondrial genes in *Ndufs4^-/-^* mice, CP2 treatment had differential effect on the AD-like and global gene expression in male and female mice. In females, global gene expression profile resembled the profile of WT female mice while CP2 treated males had distinctive profile that did not revert to the WT males. In *Ndufs4^-/-^* females, CP2 treatment rescued gene expression in coexpression Clusters B and C while in males it did not have profound effect on any of the Clusters. However, this was observed after only three days of CP2 treatment. CP2 induced changes in *Ndufs4^-/-^* females in pathways related to synaptic function, inflammation, and redox state, consistent with the previous demonstration that continuous CP2 treatment reversed transcriptomic changes in the brain tissue of APP/PS1 female mice shared with female AD patients^40^. In males, pathways affected by CP2 related to protein translation (upregulated) and RNA processing (downregulated). Furthermore, some of the transcriptomic changes associated with *Ndufs4* knockout were reinforced by CP2 even further instead of being reversed. These sex-specific response to CP2 treatment may partially be explained by the lesser disease severity, as evidenced by the concordantly downregulated genes overlapped with AMP-AD Cluster C in female but not male mice, and better therapeutic outcomes in females compared to males. These data demonstrate that inhibition of complex I activity is sufficient to induce AD-like transcriptomic signatures in the brain, but restoration of genes related to mitochondrial homeostasis and OXPHOS may not be sufficient to rescue these changes, especially at the later stages of the disease, and/or might require a longer treatment window.

Despite these findings, continuous CP2 treatment through life did not extend life span in *Ndufs4^-/-^* mice. The lack of CP2 efficacy could be explained by a short lifespan of *Ndufs4^-/-^*mice (∼ 3 months) where the window of therapeutic opportunity might have been missed. Earlier, we showed that application of CP2 to APP/PS mice starting *in utero* significantly delayed the onset of AD-like phenotype^43^. It is feasible that *in utero* application of CP2 in *Ndufs4^-/-^* mice is required to observe therapeutic efficacy in these short-lived mice. However, it is important to note that CP2 treatment did not cause toxicity or side effects and did not negatively affect weight gain or life span.

Nevertheless, the outcomes of this study did not provide unambiguous answer to whether CP2 exerts its neuroprotective properties exclusively via inhibition of complex I. Our most recent data using Cryo-EM unambiguously confirmed binding of CP2 to the isolated ovine complex I (unpublished data). The detailed investigation of the impact of *Ndufs4* knockout on the assembly and stability of complex I revealed that other subunits or OXPHOS complexes, including *Ndufaf2* and complex III, could function as stabilizing factors to explain a residual activity of complex I (∼26% in isolated mitochondria) in the *Ndufs4*^-/-^ mouse brain^30,54^. Extensive studies conducted to understand the mechanism of CP2 suggest that binding and mild inhibition of complex I is required to alter AMP/ATP ratio, activate AMPK and the subsequent signaling cascade^37–41,43^. Our studies in aging WT or symptomatic AD mice suggest that mild inhibition of complex I could be beneficial despite a presence of mitochondria with altered dynamics and function, similar to data generated with metformin^59,63^. The demonstration that mild pharmacological inhibition of complex I in *Ndufs4*^-/-^ mice reverses transcriptomics signatures of AD provides further evidence that the residual activity of complex I is required and could be sufficient to activate protective mechanisms. However, we can’t exclude the possibility of other molecular targets. Further studies will address this question.

The precise mechanisms behind sex-specific response to reduced activity of complex I and to CP2 treatment are unclear. Recent studies demonstrated that mitochondria in female rodent brain have significantly greater functional capacities, NADH-linked respiration, electron transport chain activity and ATP production compared to male brain mitochondria, which was also observed in postmortem human brain tissue^64^. The preferential utilization of lipids by female mitochondria compared to amino acids by male organelles could explain not only higher oxidative capacity of female mitochondria but also differential mechanisms associated with bioenergetic challenges and mechanisms of adaptation to energetic stress^31,65,66^. This could also relate to the fact that female *Ndufs4^-/-^* mice mimic early AD symptoms compared to male mice. Interestingly, neither heterozygous *Ndufs2^+/-^* nor *Ndufs4^+/-^* mice have reduced health or life span^67^ suggesting that the threshold required for symptom manifestation has not been reached and additional environmental or genetic risk factors are required to initiate the development of AD-like symptoms.

Our study provides a conceptual framework for the role of complex I in AD where a synergistic effect of reduced complex I activity caused by age, disease, or genetic makeup together with other genetic or environmental risk factors could initiate signaling cascade eventually affecting essential neuronal pathways that require the fidelity of mitochondrial function (Fig. 7). Reduced activity of complex I affects expression of all OXPHOS complexes and the TCA cycle, which impacts global mitochondrial functions, including redox homeostasis, synthesis of precursors for macromolecules (e.g., lipids, proteins, DNA and RNA), autophagy/mitophagy, cellular energy homeostasis, epigenetic modifications, and adaptive stress response^68,10^. Differential ability to engage compensatory mechanisms determines sex-specific disease trajectory. The outcomes of our study emphasize the necessity of assessing sex-specific differences related to disease mechanisms and a response to treatment to establish windows of therapeutic opportunity. These findings also underscore the complex nature of mitochondrial signaling that could be harnessed to improve human health. Data provide strong evidence for mitochondria as a primary instigator of AD development and as a putative therapeutic target.

**Fig. 7.**
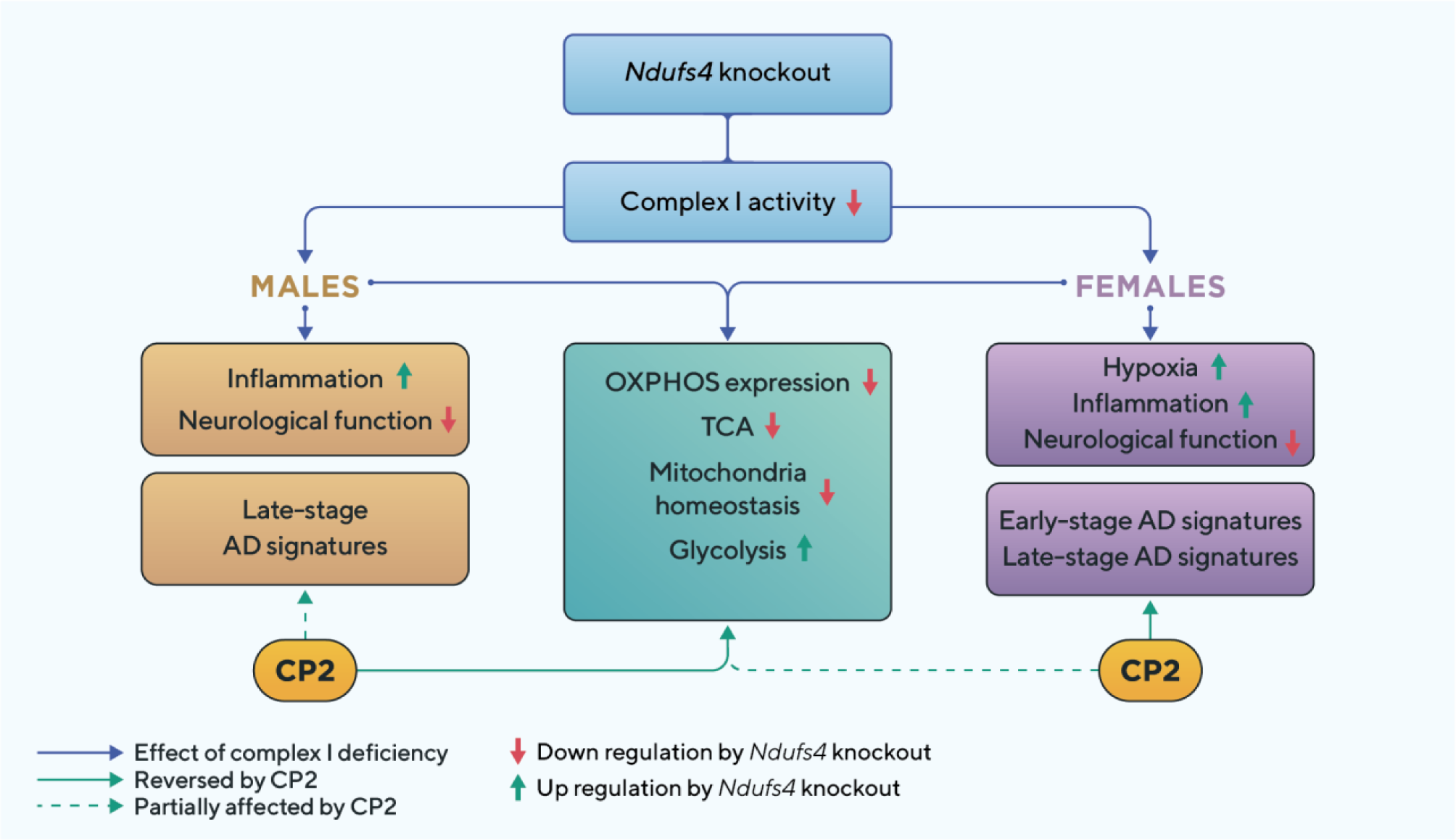
Summary of the sex-specific response of *Ndufs4^-/-^* mice to the reduced complex I activity and the development of AD-like transcriptomic signatures and to CP2 treatment.

### Study Limitations

Limitations of this study include a relatively small sample size that varied from 3 - 5 mice per group. The residual activity of complex I in brain tissue of mice studied here was not confirmed. We relied on comprehensive determination of complex I activity in the brain of *Ndufs4^-/-^* mice conducted previously^54^. The biochemical validation of transcriptomic findings remains to be conducted. Future studies should address the extent of AD-like pathology other than transcriptomic signatures in *Ndufs4*^-/-^ male and female mice. Since we did not perform functional studies in the cohorts of *Ndufs4*^-/-^ mice treated with CP2, it is unknown whether treatment had a positive effect on health span without extending life span.

## Methods

### CP2 synthesis

CP2 was synthesized by the Nanosyn, Inc biotech company (http://www.nanosyn.com) as described^69^ and purified using HPLC. Authentication was done using NMR spectra to ensure the lack of batch-to-batch variation in purity. CP2 was synthesized as free base. Stock aliquots of 20 ml were stored at −80°C. Each aliquot was used once to avoid freeze-though cycle.

### Mice and CP2 treatment

Animal study was approved by the Mayo Clinic IACUC (Protocol number A00006230-21). A whole-body homozygous *Ndufs4*^-/-^ mice^28^ were purchased from the Jackson Laboratories (B6.129S4-Ndufs4tm1.1Rpa/J, Strain #:027058). Authentication was done using PCR protocol and primers specified by the Jackson Laboratories. Breeding was conducted using heterozygous *Ndufs4*^+/-^ mice. Experiments included homozygous *Ndufs4*^-/-^ mice and WT littermates. All mice were kept on a 12 h – 12 h light-dark cycle, with a regular feeding and cage-cleaning schedule. Weight was recorded weekly. Mice were randomly selected to study groups based on their age and genotype. Continuous CP2 treatment for survival study started at 21 days of age in the cohort of male and female *Ndufs4*^-/-^ mice (*n* = 13 - 14 mice *per* group) treated with vehicle (0.1% PEG) or CP2 (25 mg/kg/day in 0.1% PEG) delivered via drinking water *ad lib* as was described previously^40,41^. Mice were treated till death. For RNA-seq analysis, *Ndufs4*^-/-^ and WT male and female mice 37-45 days of age (*n* = 3 - 5 mice *per* group) were gavaged with Vehicle (0.1% PEG) or CP2 (25 mg/kg in PEG) for 3 consecutive days. Mice were sacrificed by cervical dislocation, cortico-hippocampal region was dissected and immediately flash-frozen in liquid nitrogen.

### RNA sequencing

RNA extraction, library preparation and sequencing were performed by Novogene Inc (Sacramento, CA). Briefly, snap frozen brain tissues were homogenized using mortar grinder, and RNA was extracted using trizol based method. mRNA was polyA enriched, and quality was evaluated by Agilent BioAnalyzer, followed by ligation with Illumina Truseq adaptors, and sequenced on Novaseq 6000 (2×150 paired end, Illumina, San Diego, CA). Data in FASTQ format was quality checked by fastqc software. Low quality data and adaptor sequences were trimmed by Trimmomatic and mapped to mouse reference genome GRCm39 by STAR software with average unique mapping rate >91%. BAM files were converted to bigwig for visualization using IGV browser. Raw gene counts were then called by HTSeq excluding non-unique mapped reads. Differential expression analysis was performed using EdgeR software using *p*-values < 0.05 and log2 fold change > 0.25 using R software.

### *Ndufs4*^-/-^ genotyping

Mice was routinely genotyped using tail-clip. Briefly, the tissue was digested using Proteinase K at 50°C overnight and precipitated by saturated NaCl solution followed by centrifugation. DNA was precipitated by ice-cold EtOH, washed by 70% EtOH, air dried, and resuspended in TE buffer. DNA segments crossing the second exon of *Ndufs4* was then PCR amplified using primers as specified below and product was resolved on 2.5% TAE-Agar gel and visualized by gel-red on BioRad gel-doc imager.

Primers used:

Wild type Forward: AGT CAG CAA CAT TTT GGC AGT

Mutant Forward: AGG GGA CTG GAC TAA CAG CA

Common reverse: GAG CTT GCC TAG GAG GAG GT

### Pathway analysis and pathway-specific principal component (PC) analysis

Pathway analysis was performed using differentially expressed genes (DEGs) by EnrichR portal (https://maayanlab.cloud/Enrichr/) using Reactome2022, BioPlanet2019, WikiPathway2023, KEGG2021, Elsevier Pathway Collection, MSigDB Hallmark 2020, GO-Biological Process2023, GO-Cellular Component2023, and GO-Molecular Function2023 databases. Resultant pathways were ranked by adjusted *p*-values with top 10 presented for each figure. For pathway specific PC calculation, expression of DEGs matched to specific pathways were Z-score normalized across samples, and PC was calculated by prcomp function in R. Directionality of the PC was then examined to ensure positive alignment with the actual expression level, or it will be negatively reversed.

### Overlap with AMP-AD co-expression modules

AMP-AD co-expression modules and differential expression results were downloaded from supplemental materials of original paper^44^. Single ortholog human-mouse gene matching was used and selected mouse models were reproduced in Fig. 4a. with designated color scale.

Hypergeometric enrichment test was performed by overlapping DEGs from AMP-AD by sex and DEGs from *Ndufs4*^-/-^ mouse models, either between KO and WT or between CP2- and vehicle-treated and tested against gene universe. Concordance/discordance was calculated by first calculating concordance (KO vs WT) or discordance (CP2 vs vehicle) scores for each DE direction:

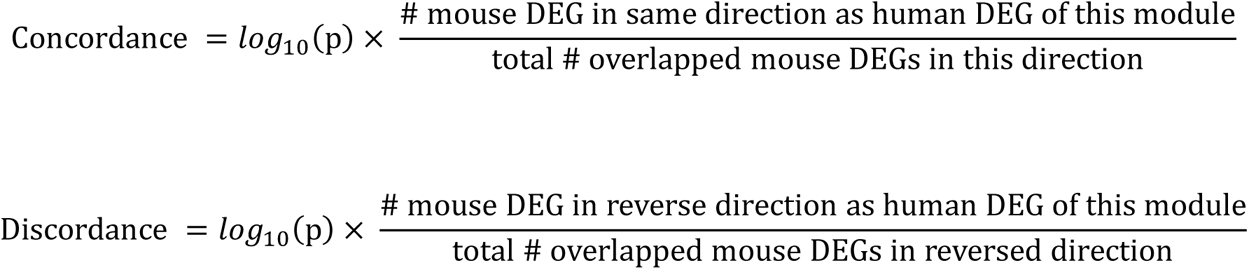

Directionalities of concordance/discordance were then determined by comparing concordance/ discordance scores of both directions using the following rules. One direction will be presented as the concordance/discordance if

1. Number of concordant/discordant gene counts in that direction is more than twice of the other direction;
2. If #1 is not satisfied, then concordance score of that direction is more than the other direction;
3. The number of concordant/discordant gene counts are more than 10. If neither direction satisfies, this module will be considered non-concordant with any mouse genes.

### Weighted gene co-expression network analysis (WGCNA)

WGCNA was performed using WGCNA package in R^70^ using all DEGs from *Ndufs4^-/-^* mice, either between KO and WT or between CP2 and vehicle-treated mice using the following parameters. Soft-thresholding power 10, minimal module size 30, deepSplit 2, merging threshold 0.2. A total of 17 co-expression modules were built using these parameters. Module cluster dendrogram was presented in Supplementary Fig. 7. Eigengenes were calculated using module Eigengenes function in WGCNA package.

## Data Availability

RNA-seq data generated in this study using mice that support the findings are deposited to public repository and are freely available at Gene Expression Omnibus with accession number GSE242286.

## Acknowledgements

This research was supported by the grants from the National Institutes of Health grant numbers RF1AG55549, RO1AG062135, AG59093 (all to ET). Its contents are solely the responsibility of the authors and do not necessarily represent the official view of the NIH and other funding organizations. The funders had no role in study design, data collection and analysis, decision to publish, or preparation of the manuscript. The authors thank Drs. R. Swerdlow, J. Wiley, J. Nguyen, H. Lee, and Mr. N. Keller for constructive suggestions and editing of the manuscript, Ms. S. Gochnauer for help with the manuscript submission, and Mr. Ivan Trushin from Trushin Studio for help with creating Figure 7.

## Author Contributions

E.T. conceived the study, assembled the multidisciplinary team of collaborators, and received funding for the project. K.J., J.N., M.O. conducted experiments; H.G. conducted RNA-seq data analysis; P.B. conducted flux analysis; C.F. participated in data analysis and interpretation. H.G. and E.T. wrote the manuscript. All authors edited the manuscript and approved its publication.

## Competing Interests

Dr. Trushina and Mayo Clinic hold three US patents on novel small molecule mitochondria targeted compounds. All authors declare no competing interests.

**Materials and Correspondence** should be directed to Dr. Trushina

## Supplementary Figure Legends

**Supplementary Fig. 1. Bigwig plot of *Ndufs4* gene expression in cortico-hippocampal tissue shows a complete knockout of exon 2 in mice subjected to the RNA-seq analysis.** *Ndufs4* exon structure is aligned at the bottom. Arrows indicate loss of exon 2 in *Ndufs4*^-/-^ mice. Tracks are normalized on y-axis and colored by genotypes. Purple: *Ndufs4*^-/-^; green: WT. Bigwig was generated using IGV.

**Supplementary Fig. 2. Pathway analysis of DEGs in brain tissue of male and female *Ndufs4^-/-^* and WT mice using the Enrichr database.** The x-axis represents negative log *p*-values of pathway enrichment; the color ramp represents percentage of genes matched for a specific pathway.

**Supplementary Fig. 3. Expression of mtDNA-encoded OXPHOS genes in male and female *Ndufs4*^-/-^ and WT mice.** Statistical differences between *Ndufs4*^-/-^ and WT mice for each gene by sex (F, Female, M, Male) were established using EdgeR package (GLM model); *p*-values are as marked.

**Supplementary Fig. 4. CP2 treatment starting at 21 days of age through life did not affect weight changes in male and female *Ndufs4^-/-^* mice.** N = 14 females treated with vehicle; n = 13 females treated with CP2; n = 14 males treated with vehicle; n = 13 males treated with CP2.

**Supplementary Fig. 5. Analysis of DEGs in *Ndufs4*^-/-^ mice treated with CP2 vs vehicle.** a-d Volcano plots of DEGs between CP2 and vehicle treated (**a**) WT female, (**b**) WT male, (**c**) *Ndufs4*^-/-^ female and (**d**) *Ndufs4*^-/-^ male mice. The x-axis represents log2 fold-change between CP2 and Veh treatments, and the y-axis represents negative log10 p-values. Counts of significant DEGs are marked on top and colored based on directionality: red, upregulated; blue, downregulated. **e** Venn diagram showing DEGs between CP2 and vehicle treated samples from *Ndufs4*^-/-^ mice overlapped between up- and down-regulated genes for female and male mice. **f**-**g** First PC of sample-wise z-score transformed expression for specific pathway colored by mouse genotype (red, vehicle; blue, CP2. The directionality of PCs was adjusted to reflect the expression level. Each dot represents one sample. Statistical difference between groups was tested using two-tailed Student’s t-test with p-values marked in the plots. **f** DEGs from assembly factors of OXPHOS. **g** PCs of all mitochondrial genome-encoded genes. **h** The expression of mitochondria genome-encoded OXPHOS genes compared between CP2 and vehicle treated *Ndufs4*^-/-^ mice. Each dot represents one sample. Red, vehicle; blue, CP2. Statistical difference between groups was tested with GLM model using EdgeR, and p-values are marked in the plots. For all boxplots, boxes represent IQR and whiskers represent 1.5 × IQR. N = 3 for *Ndufs4*^-/-^ and WT male vehicle treated groups. N = 5 for all female groups.

**Supplementary Fig. 6.** Pathway analysis of DEGs between CP2 and vehicle treated *Ndufs4^-/-^*mice by sex using the Enrichr database. The x-axis represents negative log *p*-values of pathway enrichment; the color ramp represents the percentage of genes matched for a specific pathway.

**Supplementary Fig. 7. The WGCNA cluster dendrogram**

**Supplementary Fig. 8. The eigengene plots for selected WGCNA coexpression modules significantly different between at least one group of comparison as shown in** Fig. 3a.

Eigengenes were calculated using WGCNA package, colored by treatment (red, vehicle; blue, CP2), and statistically tested between vehicle treated *Ndufs4^-/-^* and WT and between CP2 and vehicle treated *Ndufs4^-/-^* mice. Statistical difference between groups was tested using two-tailed Student’s *t*-test with *p*-values labeled in each eigengene plot. Boxes represent IQR and whiskers represent 1.5 × IQR. *N* = 3 for *Ndufs4^-/-^* male vehicle treated group. *N* = 5 for all other groups. The eigengene plots were grouped by 1) relatively up- or down-trend between vehicle treated *Ndufs4^-/-^*and WT mice; and 2) whether the difference was significant between vehicle treated *Ndufs4^-/-^* and WT mice only (*Ndufs4^-/-^* vs WT only), CP2 vs vehicle treated *Ndufs4^-/-^* mice only (CP2 only), or both and reversed directionality (rescued).

## Supplementary Tables

**Supplementary Table S1, DEG. Ndufs4 vs WT**

**Supplementary Table S2 Pathway enrichment for DEG. NdufS4 vs WT**

**Supplementary Table S3, DEG. CP2 vs Veh**

**Supplementary Table S4 Pathway enrichment for DEG. CP2 vs Veh in Ndufs4-KO mouse**

**Supplementary Table S5 WGCNA Genes by Module**

**Supplementary Table S6 WGCNA pathway analysis by module**

**Supplementary Table S7 Overlap between Ndufs4-KO mouse DEGs and AMP-AD aggregated clusters**

**Supplementary Table S8 Overlap between Ndufs4-KO mouse WGCNA coexpression modules and and AMP-AD aggregated clusters with enrichment**

